# The Genetic and Evolutionary Basis of Gene Expression Variation in East Africans

**DOI:** 10.1101/2022.02.16.480765

**Authors:** Derek E. Kelly, Shweta Ramdas, Rong Ma, Renata A. Rawlings-Goss, Gregory R. Grant, Alessia Ranciaro, Jibril B. Hirbo, William Beggs, Meredith Yeager, Stephen Chanock, Thomas B. Nyambo, Sabah A Omar, Dawit Wolde Meskel, Gurja Belay, Hongzhe Li, Christopher D. Brown, Sarah A. Tishkoff

## Abstract

**Background:** Mapping of quantitative trait loci (QTL) associated with molecular phenotypes is a powerful approach for identifying the genes and molecular mechanisms underlying human traits and diseases. How the genetic architecture of molecular traits varies across human populations, however, has been less explored. To better understand the genetics of gene regulation in East Africans, we perform expression and splicing QTL mapping in whole blood from a cohort of 162 diverse Africans from Ethiopia and Tanzania. We assess replication of these QTLs in cohorts of predominantly European ancestry and identify candidate genes under selection in human populations.

**Results:** We find the gene regulatory architecture of African and non-African populations is broadly shared, though there is a considerable amount of variation at individual loci across populations. Comparing our analyses to an equivalently sized cohort of European Americans, we find that QTL mapping in Africans improves the detection of expression QTLs and fine mapping of causal variation. Integrating our QTL scans with signatures of selection, we find several genes related to immunity and metabolism that are highly differentiated between Africans and non-Africans, as well as a gene associated with pigmentation, *TMEM216*, with evidence of population-specific selection in Nilo-Saharan speaking pastoralists.

**Conclusion:** Extending QTL-mapping studies beyond groups of European ancestry, particularly to diverse indigenous populations, is vital for a complete understanding of the genetic architecture of human traits and can reveal novel functional variation underlying human traits and disease.

## Background

Gene regulation is a principal mechanism by which genetic variation contributes to phenotypic variation, making its study essential for understanding human evolution and disease. Nearly a half century ago, King and Wilson noted the high degree of conservation between the coding regions of humans and chimpanzees, positing that non-coding variation and its effect on gene regulation must account for much of the phenotypic divergence between these species [1]. The genomics era has further underscored the importance of noncoding variation in human disease and evolution: ∼90% of the genotype-phenotype associations identified by genome-wide association studies (GWAS) cannot be explained by coding variation [2, 3], and similarly, genomic regions harboring evidence of selection in humans are significantly more enriched for variants altering expression than protein coding [4].

While GWAS and scans of selection can identify genomic regions of interest, they often lack the resolution to identify the specific genes underlying traits or targeted by selection. To bridge this gap, studies have aimed to identify genetic variation associated with fine-scale, molecular phenotypes, through quantitative trait locus (QTL) mapping [5]. Combining these molecular QTL maps with GWAS through colocalization, transcriptome-wide association studies, or Mendelian randomization, continues to prove a fruitful approach for identifying genes causally linked to traits and potential drug targets. Unfortunately, there is a persistent ancestry bias in human genomics research, with nearly 80% of GWAS participants being of recent European ancestry [6, 7], as well as the majority of participants of molecular trait studies [8], greatly limiting our ability to translate findings from GWAS to diverse populations, as well as discover population-specific variation of interest [9].

Recent studies have sought to address the genomics gap between groups of European and non-European ancestry, identifying novel GWAS associations and genetic variation contributing to gene expression differences across populations [10–14]. However, most global populations continue to be understudied, particularly in sub-Saharan Africa. Africa is the birthplace of anatomically modern humans and harbors the greatest levels of human genetic diversity across continents. Africa is home to a large array of biomes and terrains, and indigenous Africans continue to practice diverse cultural and subsistence strategies. Together, these environmental pressures have driven remarkable adaptations to infectious disease [15], diet [16], and climate [11, 17], often in a population-specific manner. These adaptive variants can have important implications for human health in Africa, and elsewhere [18], and Africa is therefore vital for our understanding of human evolutionary history and health.

In this study, we probe the genetic architecture of gene regulation in whole blood from indigenous East Africans by performing expression QTL (eQTL) and splicing QTL (sQTL) mapping in a cohort of 162 individuals, representing nine ethnic groups, from Ethiopia and Tanzania. We measure the degree to which African architecture is shared with that of non-Africans, test whether Africans harbor functional variation absent from existing cohorts, and investigate the demographic and genetic forces that may contribute to variation in gene regulatory architecture. We test whether fine-mapping of QTL signals is improved in Africans relative to an equivalently sized cohort of European Americans, and highlight individual genes with improved fine-mapping in Africans. Finally, we measure the effect of selective forces on shaping gene regulatory architecture and identify candidate genes under selection.

## Results

### Population Structure

The cohort for this study consists of 171 Ethiopian and Tanzanian individuals belonging to nine ethnically and culturally diverse sub-Saharan groups, including the Cushitic speaking Agaw and Weyto, the Semitic speaking Argoba and Amhara, the Omotic speaking Dizi, the Nilo-Saharan speaking Mursi, and the Chabu who speak an unclassified language similar to Nilo-Saharan, and the Khoesan speaking Hadza and Sandawe (Figure 1A). These populations practice a variety of subsistence strategies, including foraging (Hadza and Chabu currently, Sandawe and Weyto formerly), with a diet diverse in foraged tubers, fruit, and hunted game; pastoralism (Mursi), a lifestyle that revolves around cattle herding and a diet high in animal proteins and fats; agriculturalism (Agaw, Amhara, and Argoba), a sedentary lifestyle with a diet high in cultivated carbohydrates; and agropastoralism (Dizi), which relies on both crops and livestock.

**Figure 1:**
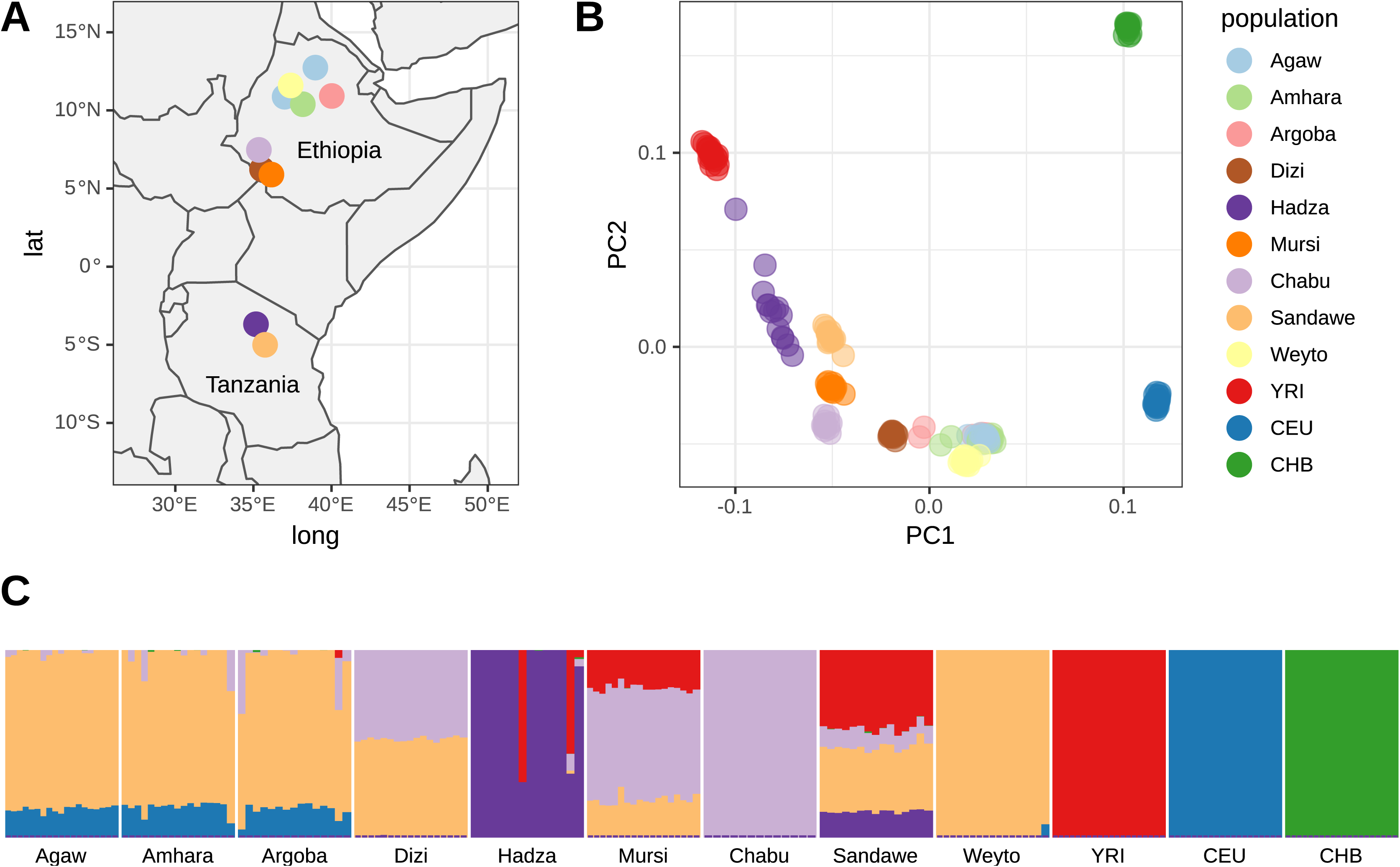
Global and genetic structure of study populations. **A)** Locations of East African populations sampled in this study across Ethiopia and Tanzania. **B)** Principal Component Analysis of genetic data across 162 East Africans, combined with 20 West African Yoruba (YRI), 20 European Americans (CEU), and 20 Han Chinese (CHB) from the 1000 Genomes Project. **C)** ADMIXTURE analysis of East African, YRI, CEU, and CHB populations.

To investigate the genetic diversity and structure of these populations, a subset of 162 individuals were genotyped at approximately 4.5 million SNPs on the Illumina Omni5M Exome array. These data were further imputed using a reference panel composed of the 1000 Genomes Project (1kGP) dataset [19] and a dataset of whole genome sequences (WGS) from 180 sub-Saharan African individuals (methods, unpublished). To place their genetic variation in a global context, genotype data from the nine study populations were merged with 1kGP WGS data from 20 individuals each of Yoruban (YRI), Northern and Western European (CEU), and Han Chinese (CHB) ancestry (methods). Principal component analysis (PCA) of this merged dataset recapitulates a primary separation between African and non-African individuals along the first PC, explaining 3.8% of the variance. The second PC, explaining 1.8% of the variance, further separates CEU and CHB individuals, as well as East Africans and the YRI (Figure 1B). Higher PCs further separate variation in Africa; PC3 captures variation between the Hadza and YRI, and PC4 between the Hadza and Chabu. Several groups cluster relatively nearer to CEU Europeans along PC1, most notably the Ethiopian Agaw, Amhara, Argoba, and Weyto, which are known to have moderate levels of Eurasian admixture [20, 21]. Inferred ancestry components from *ADMIXTURE* [22] also estimate components of non-African ancestry among these Ethiopian groups, as well as admixture with Bantu-speaking populations of Western African origin [19], represented by the YRI, in the Sandawe, Mursi, and Hadza (Figure 1C).

### Transcriptomic traits in Africans

To assess the contribution of genetic variation to transcriptomic trait variation, we performed genome-wide QTL mapping for expression (eQTL) and splicing (sQTL) transcriptomic traits in *cis* for expressed protein-coding and long-noncoding RNA genes; collectively we will refer to eQTLs and sQTLs as transcriptomic QTLs (tQTLs). We first correct our phenotypes (expression and splicing) for a number of covariates, including age, sex, delivery date, hidden covariates inferred by *PEER* [23], and cell-type fractions inferred by *CIBERSORT* [24]. Cell-type composition of whole blood is known to vary between individuals, and to be a source of confounding in QTL studies [25]. To account for ancestry and relatedness, we generate a genetic relatedness matrix (GRM) and perform tQTL mapping using the linear mixed model tool *GEMMA* [26]. Testing all autosomal SNPs with minor allele frequency (MAF) greater than 0.05 and within 100kb of the target gene transcription start site (TSS) for eQTLs or within 100kb of the target intron for sQTLs, we identify 99,685 SNPs associated with the expression of 1,330 genes (eGenes) and 74,445 SNPs associated with splicing of 1,118 introns (sIntrons) in 776 genes (sGenes) at FDR < 0.05 (methods).

SNPs associated with expression (eSNPs) and splicing (sSNPs) show a characteristic enrichment near the transcription start site or intron boundary of their target gene, respectively [27] (Figure 2A), and are enriched in a variety of functional categories, including transcription start sites, enhancers, and splice sites, and are depleted in repressed chromatin regions. We also find a significant overlap with chromatin QTLs (caQTLs) identified in lymphoblastoid cell lines (LCLs, Figure 2B). Further, alleles associated with increased chromatin accessibility are significantly more likely to be associated with increased expression (OR = 2.9, p = 8.2 x 10^-37^ Fisher’s Exact Test) and slightly less likely to be associated with increased junction inclusion (OR = 0.82, p = 0.03 Fisher’s Exact Test), suggesting that regulatory mechanisms altering chromatin accessibility play a greater role in regulation of gene expression than splicing. When we restrict to variants with a greater than 10% probability of being causal (methods), we find a further enrichment in functional categories, particularly for caQTLs among eQTLs and splice regions among sQTLs, indicating we are capturing true causal variation (Figure 2B).

**Figure 2:**
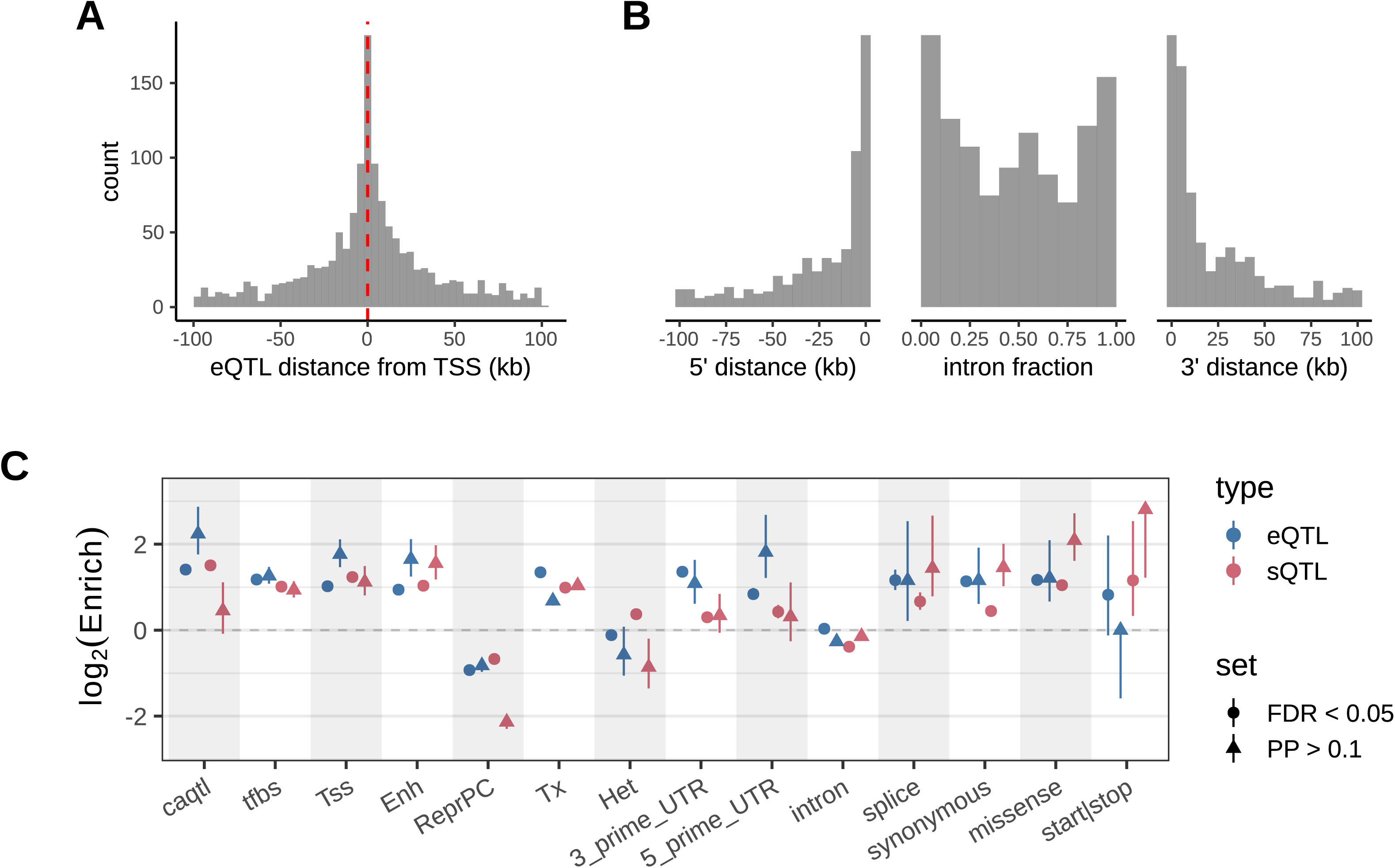
Genomic context of tQTLs. **A)** Enrichment of top eQTLs near the transcription start site (TSS) of their target gene. **B)** Enrichment of top sQTLs near the intron boundary of their target intron. Densities of sQTLs are separated depending on whether they’re upstream of the target intron (5’ distance), within the intron (intron fraction), or downstream of the intron (3’ distance). **C)** Enrichment of tQTLs across functional categories, stratified by FDR significance and posterior probability (PP) of being causal. Categories include chromatin accessibility QTLs (caQTL) in LCLs from Tehranchi *et al.* [33]; transcription factor binding sites (TFBS) for 140 transcription factors in GM12878 LCLs [34]; transcription start sites (TSS), enhancers (Enh), Polycomb-repressed chromatin (ReprPC), transcribed (Tx), and heterochromatin (Het) annotations from ChromHMM in GM12878 LCLs [34]; and 3’ UTR, 5’ UTR, intron, splice site, synonymous, missense, and start gain/loss or stop gain/loss annotations from Variant Effect Predictor (VEP) [88].

Of the genes tested, 198 have both an eQTL and sQTL in our cohort, suggesting possible shared genetic architecture between these transcriptomic traits. To evaluate whether eQTLs are enriched for sQTLs overall, we first compute the π_1_ statistic, which measures the estimated fraction of sQTLs that are true positives in the eQTL scan. A π_1_ value of 0.61 suggests that the majority of sQTLs affect expression or are in LD with variants affecting expression (Figure S3), though many of these fail to reach genome-wide significance. To further evaluate whether the genome-wide significant eQTL and sQTL signals are driven by shared causal variants, we estimated 90% credible sets for each set of QTLs, defined as the minimal set of variants which have at least a 90% probability of containing the causal variant, using the probabilities estimated above (methods). Overall we find overlapping credible sets for 114 of the genes with both a significant eQTL and sQTL, which makes up about 9% (114/1,330) of all eGenes in our cohort, comparable to the 12% overlap observed in GTEx [28]. Taken together, this observation suggests that splicing variants likely cause subtle but detectable changes in gene read counts, but that the genetic variants driving genome-wide significant eQTLs and sQTLs are largely independent.

### Replication of tQTLs in non-Africans

To validate our tQTLs, and to assess sharing of molecular trait architecture between cohorts of predominantly African vs. predominantly European ancestry, we compared our results to whole blood analyses from the Genotype-Tissue Expression project (GTEx) v8, which is comprised of 85% European Americans [28]. For those QTLs tested in both cohorts, we find that both eQTLs and sQTLs identified in the African cohort show overall high reproducibility in GTEx, with π_1_ values for eQTLs and sQTLs of 0.88 and 0.90, respectively (Figure S4, methods). In addition to π_1_, effect sizes between our cohort and GTEx also show overall strong concordance (Pearson’s = 0.73 for eQTLs and 0.82 for sQTLs, Figure 3: Replication of tQTLs between East Africans and GTEx v8). To assess whether the observed replication is significantly affected by the different genome versions used between our study and GTEx v8, we also measured π_1_ of eQTLs in GTEx v7, finding a π_1_ of 0.83 (Figure S4). Those tSNPs that fail to replicate in GTEx (p > 0.01) show consistently lower MAF (Figure 3: Replication of tQTLs between East Africans and GTEx v8); this failure to replicate includes the top eSNP in Africans for 308 genes and the top sSNP for 220 introns in 185 genes, indicating widespread differences in power for detecting tQTLs across ancestral groups.

**Figure 3:**
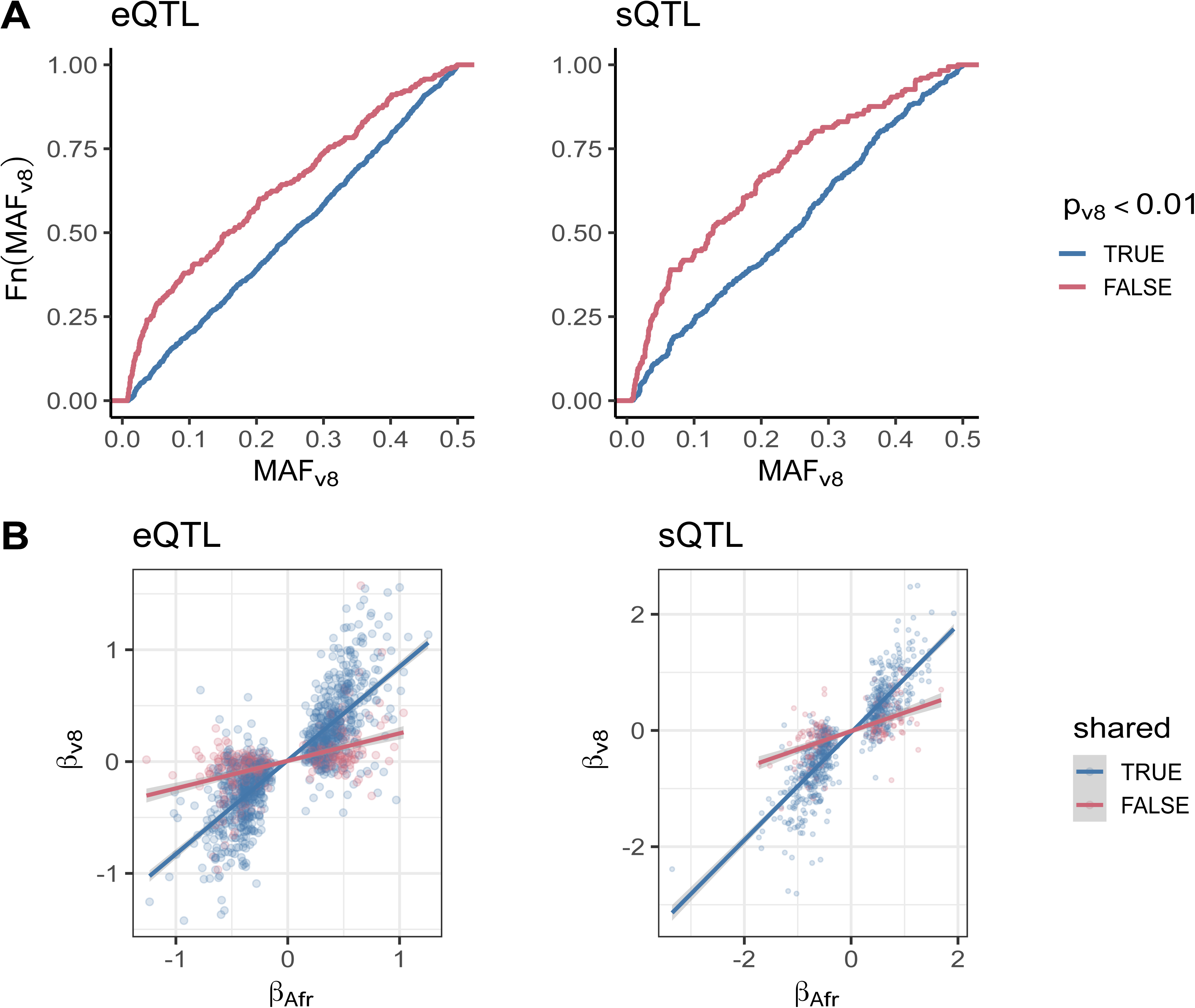
Replication of tQTLs between East Africans and GTEx v8. **A)** Minor allele frequency distribution in GTEx v8 of FDR-significant tQTLs identified in East Africans, colored by whether they have a p-value less than 0.01 in GTEx v8. **B)** Comparison effect sizes of tQTLs identified in East Africans. Lines show the best fit regression line between East Africans and GTEx v8 effect sizes, colored by whether the tQTL is shared (i.e. is no longer significant after conditioning) or is independent (remains significant after conditioning).

We next investigate whether expression differences may affect replication between cohorts. Of the 1,330 eGenes identified in Africans, the expression of 98 in GTEx v8 whole blood is too low to be tested for eQTLs. These 98 genes are significantly enriched in two KEGG pathways, “Hypertrophic cardiomyopathy” (FDR = 0.032) and “Dilated cardiomyopathy” (FDR = 0.038). Investigating what may be driving broader expression differences for testable genes, we identify those genes measured in Africans that fail to reach expression thresholds for testing in GTEx whole blood and vice versa. Altogether 951 out of 12,377 genes measured in both cohorts and tested for eQTLs in Africans were not tested in GTEx. These genes are enriched for a number of biological processes related to sensory perception, including perception of smell (FDR = 2.85 x 10^-6^), sound (FDR = 1.60 x 10^-5^), mechanical stimulus (FDR = 5.60 x 10^-5^), and chemical stimulus (FDR = 5.22 x 10^-4^). Similarly, 6,728 out of 18,168 tested for eQTLs in GTEx were not tested in Africans and are enriched for several biological processes related to immunity, including “complement activation, classical pathway” (FDR = 1.78 x 10^-22^), “humoral immune response mediated by circulating immunoglobulin” (FDR = 7.32 x 10^-18^), and “B cell mediated immunity” (FDR = 2.02 x 10^-2^). This observation suggests that disease status, sample collection, and response to environmental factors, in addition to genetics, may account in part for incongruent findings between eQTL cohorts.

While tQTLs as a whole show strong replication using π_1_, we also investigate the degree to which individual loci show evidence of shared causal variation. Estimating credible sets for all eGenes and sIntrons in GTEx v8 as described above, we find that 715/1262 (57%) of eGene credible sets and 619/852 (73%) of sIntron credible sets in Africans overlap with credible sets in GTEx v8. While the majority of tQTL credible sets overlap, the many non-overlapping sets suggests many tQTL signals identified in Africans may be driven by independent causal variants. To further evaluate this independence we remapped tQTLs in Africans, conditioning on sets of independent tQTLs identified in GTEx by forward regression [28]. In cases where there are no genome-wide significant eQTLs or sQTLs in GTEx (169 genes and 541 introns, respectively) we instead condition on the lead eSNP or sSNP in GTEx. Using the original FDR significance thresholds for calling eQTLs and sQTLs, we find that 362 (27%) of eGenes and 224 (20%) of sIntrons remain significant after conditioning on GTEx SNPs, including the top variants for 328 eGenes and 199 sIntrons, suggesting widespread independent causal variation in Africa.

Investigating what may be driving the independent signals in our cohort, we compare minor allele frequency (MAF), linkage-disequilibrium (LD) structure, and effect size differences between our cohort and GTEx v8 samples or European-ancestry proxies (CEU individuals from the 1kGP, methods). For 8 genes, *INPP5K*, *TMEM140*, *ACSM3*, *CNTNAP3*, *PPP1R14C*, *PDZK1TP1*, *GPR56*, and *TRAM2*, the top eSNP in Africans is untested in GTEx and has a MAF < 0.01 (the threshold used by GTEx) in 1kGP EUR populations. Similarly, the top sSNPs for 4 genes, *ADAM8*, *ICAM2*, *LINC00694*, and *MAPK1* are absent in GTEx and have a EUR MAF ≤ 0.01. Overall, however, we find that frequency differences between Africans and EUR are similar between shared and independent tQTLs (Figure S6). To investigate the impact of LD variation on tQTL replication, we estimate *r^2^* between tQTL lead SNPs and SNPs within 100kb of lead SNPs in 1kGP CEU and YRI populations. We find that correlations between CEU and YRI *r^2^* values do not differ significantly between shared and independent tQTLs (Figure S6). Finally, comparing effect size variation, we find a significant reduction in effect size correlation between Africans and GTEx among independent tQTLs relative to shared signals (Figure 3: Replication of tQTLs between East Africans and GTEx v8, p < 2.2 x 10^-16^), which may reflect true effect size variation, GxE effects [13,14,29], or possibly more subtle differences in MAF and local LD between these cohorts [30].

### Fine Mapping

In addition to assessing the replication of transcriptional QTLs in the larger GTEx v8 dataset, we are interested in the relative power to detect and fine-map tQTLs between cohorts of predominantly African versus European ancestry. To account for sample size differences between our cohort and GTEx, we performed eQTL mapping in a size-matched sample of 162 European-American (EA) individuals from GTEx v8 using *FastQTL* [31], with sex, sequencing platform, PCR batch, the top 15 *PEER* factors, and top 5 genotype PCs as covariates. Testing all SNPs with MAF > 0.05 within 100kb of the target TSS, we identify 1,029 eGenes in the 162 EA individuals at FDR < 0.05, compared with 1,330 identified in Africans, of which 326 eGenes are FDR-significant in both cohorts. Despite only 326 eGenes being shared, we find consistently high replication in an independent whole blood meta-analysis [32]; eQTLs that are FDR-significant in both cohorts reach a π_1_ of 0.999, while eQTLs discovered only in Africans reach a π_1_ of 0.958 and eQTLs discovered only in EAs reach a π_1_ of 0.989. This observation suggests that the greater number of eGenes in Africans is not driven by an increase in false positives, and that at similar sample sizes, Africans have an improved power to detect eQTLs compared with individuals of European ancestry.

We next investigate the relative ability to fine-map eQTLs between our African cohort and the 162 EA individuals from GTEx v8. Considering eGenes that are FDR-significant in either cohort (methods), we perform fine-mapping in both our African cohort and the 162 EAs using the approach described above. Overall, most genes do not fine-map well at this modest sample size, with 57% of genes having a credible set larger than 50 in both cohorts (Figure 4: Fine mapping in East Africans vs. GTEx v8). Excluding these genes, we find that Africans have a smaller credible set in 63% of cases (437/697, p = 2.06 x 10^-11^ binomial test), with a median credible set size of 25 in Africans vs 58 in EAs, and 23 genes fine-mapped to a single variant in Africans vs. 13 in EAs. One possible explanation of the smaller credible sets in Africans is that Africans simply have fewer SNPs tested per gene; however, we find the opposite, with 94% of genes have fewer tested SNPs in EAs.

**Figure 4:**
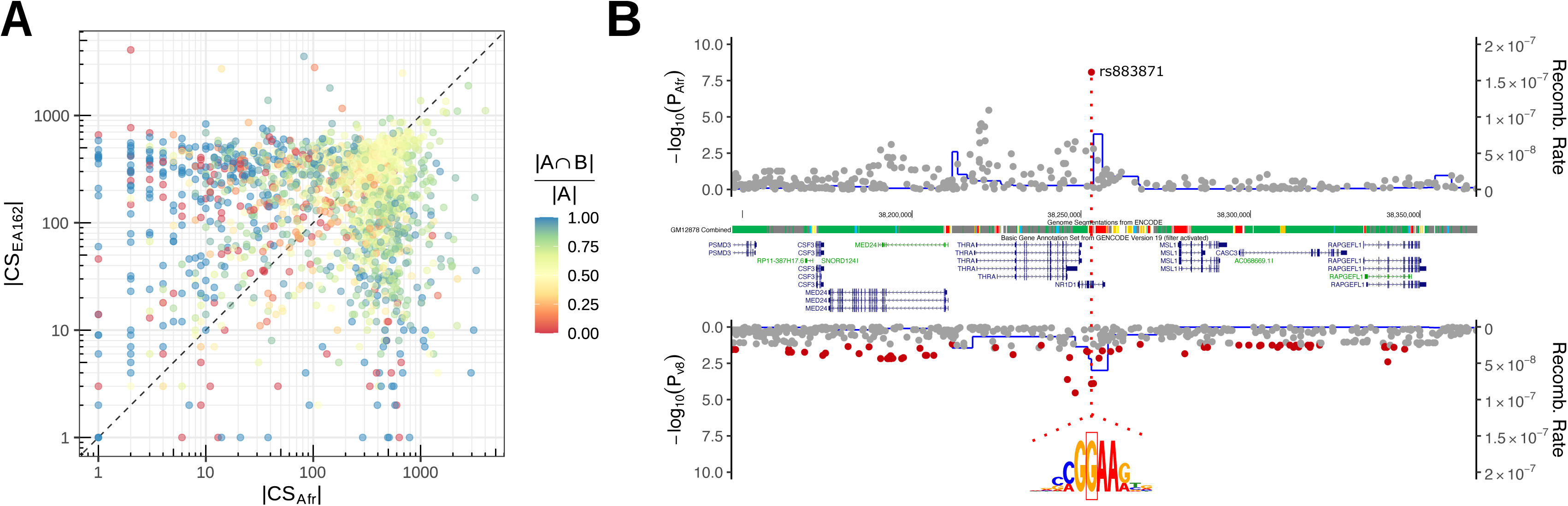
Fine mapping in East Africans vs. GTEx v8. **A)** Credible set (CS) sizes for eGenes identified in 162 East Africans (Afr) or a subset of 162 European Americans from GTEx v8 (EA162). Points are colored by the fraction of SNPs in the smaller credible set A that are shared with the larger set B, 1 indicating that the smaller set is a subset of the larger set, and 0 indicating the smaller set shares no SNPs with the larger set. **B)** Locus plot of *NR1D1* eQTLs identified in 162 East Africans (Afr) or the full GTEx v8 cohort (v8). P-values are overlaid with African (YRI) and European-American (CEU) recombination rates, GENCODE v19 [81] gene models from the UCSC genome browser [94] (http://genome.ucsc.edu) and inferred ChromHMM[95] states for GM12878 [34]. The top SNP in Africans, rs883871, disrupts a nucleotide for the core motif of ETS-family transcription factors (motif of *ETS1* shown).

We further compare our credible sets in African eQTLs to credible sets estimated in the full GTEx dataset. As expected, the majority of genes have smaller credible sets in GTEx due to the considerably larger sample size (670 vs 162), though we do identify several examples of greatly reduced credible sets in the African cohort. For 18 eGenes and 32 sGenes we are able to fine-map the QTL signals to a single variant in Africans and find that these variants overlap a lead GWAS association for 10 eGenes and 3 sGenes (supplement). We highlight rs883871 (Figure 4: Fine mapping in East Africans vs. GTEx v8), an eQTL for both *THRA* and *NR1B1*, which is FDR-significant in GTEx whole blood but is not the lead eSNP. rs883871 is a strong chromatin QTL in lymphoblastoid cell lines (LCLs) [33], overlaps the binding sites of numerous transcription factors (TFs) in the LCL GM12787 [34], is predicted to disrupt a consensus motif for the ETS family of TFs, which share a core ‘CCGGAA’ motif, and is the lead SNP for a Multiple Sclerosis GWAS association [35]; variants in *ETS1* itself have been previously associated with Multiple Sclerosis [36]. Given our modest sample size compared with GTEx, we expect that mapping of tQTLs and other molecular traits in larger cohorts of genetically diverse populations will further enhance fine-mapping of QTLs, and when combined with more diverse GWAS studies, may identify novel causal genes underlying human traits and disease.

### Signatures of Selection

Gene regulation is known or suspected to underlie many adaptive traits in humans, including diet [16, 37], immunity [38], and skin pigmentation [11], and transcriptomic traits show evidence of both purifying and positive selection [13,14,39]. Consistent with previous tQTL studies we find decreasing effect size with increasing MAF among eQTLs and sQTLs, indicative of negative selection against variants of large effects (Figure S7). To identify QTLs with evidence of positive selection we measure genome-wide *F_ST_* between our broader African dataset and the 1kGP European (EUR) individuals, with the expectation that selection for expression-altering alleles will lead to increased differentiation at these loci. To assess whether tQTLs are enriched for evidence of positive selection we identify the highest *F_ST_* value for all SNPs in high LD (*r^2^* > 0.8) with the top eQTL or sQTL and compare these values with null SNPs matched on MAF and the number of SNPs in LD (methods). Overall, we do not find an enrichment of high *F_ST_* among eQTLs or sQTLs, suggesting that selection has not driven significant frequency differentiation at the majority of tQTLs (Figure S8).

We next investigate evidence of selection at individual loci. To account for the fact that the top eSNP may not be the true causal SNP, we score an individual gene’s evidence of selection by taking a weighted sum of each SNP’s *F_ST_* value multiplied by the probability of that SNP being causal (methods). Considering loci with a score within the 99^th^ percentile threshold of all SNP *F-_ST_* values as candidates, we identify 27 eGenes and 25 sGenes with evidence of selection (supplement). The most differentiated eGene is *TTC26* (weighted *F_ST_* = 0.59); a mutation in this gene has been associated with abnormal cilia in model organisms and biliary ciliopathy in human liver [40]. We also identified a strong signature of selection at *TMEM154* (weighted *F_ST_* = 0.59, Figure 5A), a mostly uncharacterized gene that has been associated with Type II Diabetes Mellitus and beta cell function in humans and lentiviral infection in sheep [41, 42]. Other highly differentiated loci include Platelet Factor 4 Variant 1 (*PF4V1*, *F_ST_* = 0.50), *IL8* (*F_ST_* = 0.49), a major inductor of immune cell chemotaxis and activation [43], and *CCR1* (*F_ST_* = 0.43), a chemokine receptor. Among the most differentiated sGenes we find several related to immunity and metabolism, including *NADSYN1* (weighted *F_ST_* = 0.50), a gene associated with vitamin D concentration [44], *BTN3A3* (weighted *F_ST_* = 0.50), a butyrophilin gene implicated in activation of T cells [45], and *GANC* (weighted *F_ST_* = 0.43), a member of the glycosyl hydrolase family 31, which play a key role in glycogen metabolism [46].

**Figure 5:**
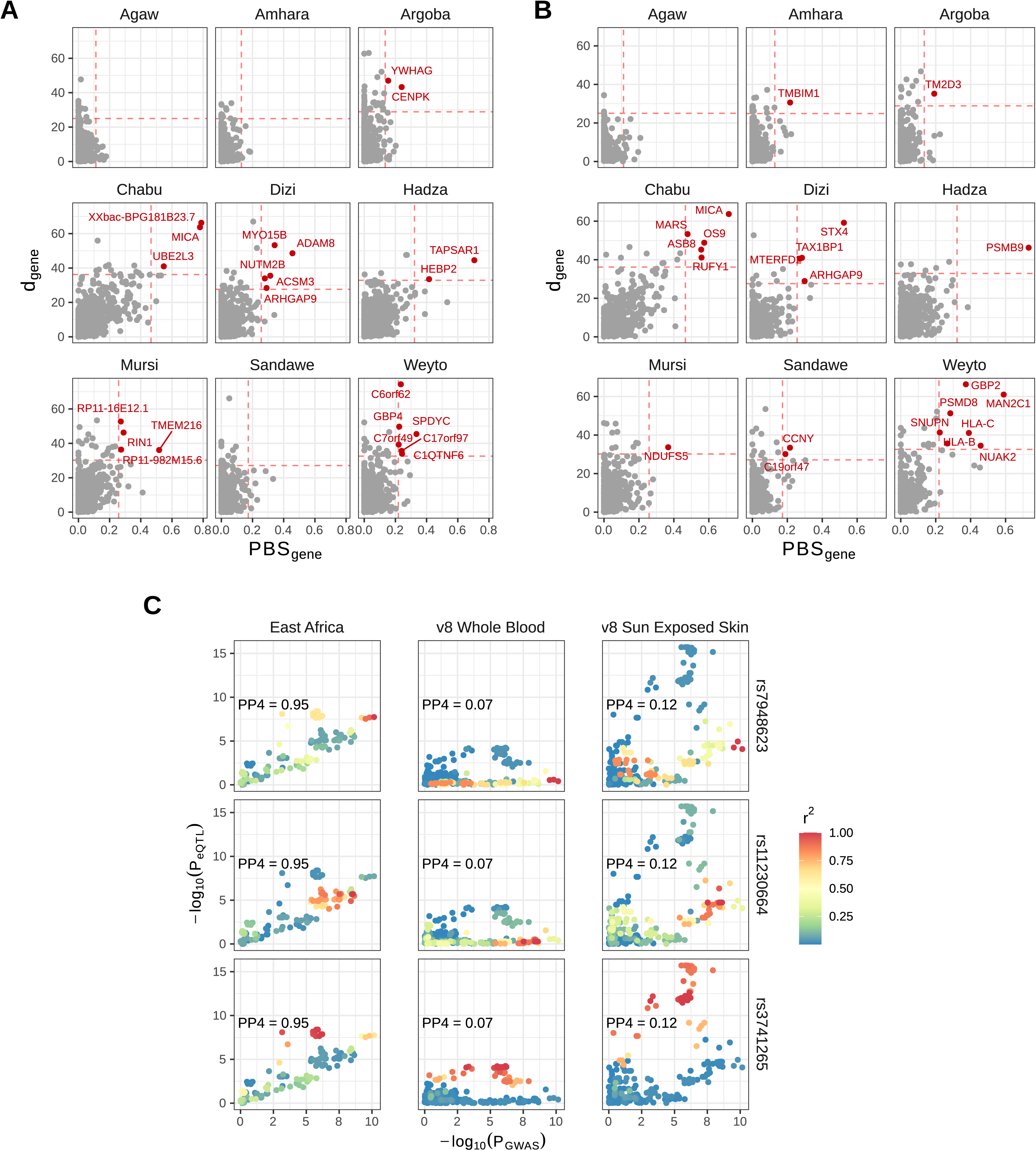
Population-specific selection in East Africa. **A)** Gene scores for the *d*-statistics plotted against the population branch statistics (*PBS*) for each population. *PBS* is calculated for each focal population versus the CEU and YRI populations from the 1000 Genomes Project. Genes with a score above the 99.5^th^ percentile of genome-wide statistics for *d* and *PBS* are highlighted in red. **B)** Comparison of pigmentation GWAS p-values from Crawford *et al*. [11] against eQTL p-values from our study (East Africa), GTEx v8 Whole Blood, or GTEx v8 Sun-exposed skin (lower leg), in the style of LocusCompare [96]. Variants are colored by their degree of LD with three top pigmentation GWAS variants, rs7948623, rs11230664, and rs2512809. Colocalization probabilities from *coloc* [86] (PP4) are indicated for each eQTL group.

Given our genetically and culturally diverse cohort we are also interested in tQTLs with evidence of population-specific differentiation and selection. For each of the nine populations in the African dataset we calculate a modified version of the *d* statistic [47], a summation of normalized, pairwise *F_ST_*, which tests for variants that are highly differentiated in a focal population versus other populations (methods). As above, we weight these *d*-statistics by the probability of a SNP being causal to derive a ’*d*-score’ for each gene or intron. Genes with high *d-*scores in populations with evidence of non-African admixture (i.e. Agaw, Amhara, Argoba, and Weyto) are more genetically similar to EUR samples from the 1kGP, based on *F_ST_*. Conversely, populations with evidence of west-African admixture (i.e. the Hadza, Mursi, and Sandawe) are more genetically similar to YRI samples at high *d*-score genes, suggesting that in many cases the genetic differentiation at these loci is driven by population-specific patterns of admixture. We therefore calculate the population branch statistic between (*PBS*) [48] between individual populations in our study and 1kGP CEU and YRI populations. Considering genes with a weighted *d* and *PBS* score in the top 99.5^th^ percentile as significant, we identify 22 eGenes and 22 sGenes with significant evidence of population-specific selection (Figure 5: Population-specific selection in East Africa. and B).

Among the top eGenes with evidence of population-specific selection is *TMEM216* among the Nilo-Saharan speaking Mursi pastoralists (Figure S9). This gene is located near a skin pigmentation GWAS locus discovered in a cohort with the same sub-Saharan African populations [11]. This association signal overlaps the UV-repair gene *DDB1*, as well as several other genes expressed in melanocytes. Colocalization analyses show strong overlap between the African *TMEM216* eQTL and pigmentation GWAS signals (PP4 = 0.95, Figure 5C, methods), suggesting possible shared causal variation between *TMEM216* expression and pigmentation variation. LD patterns around *TMEM216* shows evidence of three independent eQTLs segregating for this gene, tagged by rs7948623, rs11230664, and rs3741265. Two of these SNPs, rs7948623, rs11230664, are also genome-wide significant GWAS SNPs for pigmentation variation in Africans, while the third, rs3741265, is marginally significant (p < 10^-5^, Figure 5C), All three SNPs show strong population-specific differentiation in Ethiopian Nilo-Saharan groups, who have amongst the highest levels of skin melanin of any global population (Figure S9). Previous analyses of these populations have shown evidence of a selective sweep near this pigmentation GWAS locus, including high *PBS* and *d* values among GWAS variants (Figure S10) and extreme negative Tajima’s D values overlapping the *TMEM138/TMEM216* locus [11].

The top GWAS variant, rs7948623, overlaps an active enhancer in keratinocytes and melanocytes and has been demonstrated to alter enhancer activity in melanocytes via luciferase reporter assays [11]. rs7948623 is a significant eQTL for *TMEM216* in our study but is not significant in GTEx whole blood, though it is in ovary, nerve, and exposed skin. In addition, rs7948623 is a significant sQTL for *TMEM216* in multiple GTEx tissues, including exposed skin (Figure 5C). A second group of *TMEM216* eQTL and pigmentation GWAS variants are tagged by rs11230664 and include the indel rs148172827, which overlaps an active melanocyte enhancer, and shows significant correlation with *TMEM216* expression in GTEx exposed skin (Figure 5C). We do not identify significant sQTLs in Africans for *TMEM216*, however the top sSNP for *TMEM216* in GTEx exposed skin, rs3741265 (p = 1.43 x 10^-322^), is in high LD with the top TMEM16 eQTL in Africans, rs7934229 (*r^2^* = 0.99). Both of these SNPs are moderately associated with skin pigmentation in Africans (p < 5 x 10^-6^) but do not reach genome-wide significance (Figure S11).

## Discussion

This study extends our understanding of the genetic basis of human gene regulation, with the inclusion of whole blood samples for 162 ethnically diverse sub-Saharan Africans from Ethiopia and Tanzania. We find that variation underlying expression and splicing is broadly shared between African and European cohorts, though there is considerable independent variation at individual loci in Africans, often driven by variation in frequency and effect sizes of tQTLs. When matched for sample size, Africans show improved fine mapping of molecular traits, facilitating the identification of causal variants and candidate genes underlying GWAS traits. This diverse cohort also allows for inference of tQTLs with evidence of local adaptation, identifying *TMEM216* as a target of selection in Nilo-Saharan speakers and a candidate gene that may play a role in skin pigmentation.

We find that the majority of tQTLs replicate between Africans and GTEx v8, with π_1_ values near 0.9 among both eQTLs and sQTLs, on par with the 0.919 value estimated between African Americans in the GENOA cohort [49] and EUR populations from the Geuvadis project [12]. We also observe strong effect size correlation between tQTLs in our study and GTEx v8.

Investigating individual loci, however, we find that many genome-wide signals are driven by distinct causal variation; 43% of eQTL and 27% of sQTL credible sets in Africans do not overlap those in GTEx v8, and 27% of eGenes and 20% of sIntrons have QTL signals that remain significant after conditioning on all tGTLs in GTEx.

Investigating what may account for QTL differences between Africans and non-Africans, we find that genes relating to sensory perception and immunity show differential expression between our African cohort and the GTEx cohorts, pathways known to vary across populations and environments [50, 51]. Additionally, the post-mortem nature of GTEx samples may contribute to expression differences. An analysis of the effects of death on gene expression in GTEx found that immune genes in whole blood are significantly dysregulated following death, however this change was characterized by an overall deactivation of immune genes, along with an overall increase in NK cells and CD8 T-cells and a reduction in neutrophils [52]. In addition to expression differences, we find an enrichment for low frequency variants in GTEx among non-replicating tQTLs. However, the majority of tQTLs that are conditionally independent show similar frequency differences with shared tQTLs, suggesting that frequency variation alone cannot account for independent tQTLs. This issue of trans-ethnic GWAS replication is an ongoing area of research [53, 54], and non-replication may occur for many reasons including frequency variation, differences in power, LD, or true differences in effect size, including G x E effects. While we do not find a significant difference in local LD structure between shared and independent QTL signals, we do find significant differences in estimated effect sizes. Using a Bayesian approach to account for frequency and LD variation, Brown *et al*. also found eQTL effect size differences between EUR and YRI individuals from Geuvadis [12], which become more pronounced as genetic effects become weaker [55]. However for strong, genome-wide significant effects, Zanetti and Weale demonstrated using simulations that most trans-ethnic differences in GWAS effect sizes can largely be accounted for by a combination of frequency and LD variation, though they could not rule out effect size differences [30].

Beyond replication, we demonstrate that at comparable sample sizes, African cohorts have improved sensitivity to detect tQTLs and improved ability to fine-map causal variants, compared with cohorts of European ancestry. It is well established that non-African populations have more extensive LD relative to Africans [56, 57], resulting from the out-of-Africa bottleneck [58, 59], which likely accounts for the observed improvement in fine-mapping in African populations. As to the increased sensitivity to detect tQTLs in Africans, one hypothesis is a higher false-positive rate in the African cohort. However we find comparable replication of African-specific tQTLs in a large, independent meta-analysis [32], suggesting that false positives do not account for the observed improvement. Moreover, Quach *et al.* found a similar pattern of improved sensitivity to detect eQTLs in individuals of self-reported African ancestry in an analysis of stimulated and unstimulated monocytes from 200 Belgians, 100 of European and 100 of African ancestry [60]. Among African Belgians they found 13% more eQTLs in unstimulated monocytes, and 10% more eQTLs across all conditions. While several other studies have mapped eQTLs across multiple ancestry groups [12,14,61,62], variation in sample size precludes direct comparison of sensitivities across ethnicities.

In addition to the inclusion in our study of ancestral groups not represented in existing reference cohorts (e.g. the 1kGP), which enables the detection of novel regulatory variation, these populations live in diverse climates and have distinct cultural and subsistence practices, which may have driven unique local adaptations. Using an outlier approach based on the *F_ST_* based *d* and *PBS* statistics [47, 48], we identify population-specific differentiation of tQTLs among East African populations. One notable example is the eQTL *TMEM216* among the Mursi, which is near a recently identified pigmentation locus specific to sub-Saharan Africans [11]. *TMEM216,* and the nearby *TMEM138* gene, form an evolutionarily conserved *cis*-regulatory module vital for ciliogenesis, and have been identified as causal genes underlying Joubert and Merkel syndromes [63, 64]. *TMEM216* has not been previously associated with pigmentation variation, though activation and suppression of primary cilia have been shown to inhibit and activate melanogenesis, respectively, in a human skin model [65]. Consistent with this, we find that the expression decreasing allele is associated with increased melanin levels for rs7948623, rs11230664, and rs3741265, and is most common in the Mursi, a populations with darkly pigmented skin (Figure S9)[11]. In addition, recurrent somatic mutations driving alternative splicing of *TMEM216* are significantly associated with melanoma in The Cancer Genome Atlas (TCGA), suggesting possible tumor suppressor function for this gene [66]. While the strong colocalization between the *TMEM216* eQTL and pigmentation GWAS signals suggests *TMEM216 as* a possible pigmentation gene, there are several haplotypes segregating in this region, some of which carry tQTLs for other genes in GTEx (Figures S12 and S13). In addition, several nearby genes show melanocyte-specific expression, or have been previously associated with pigmentation in other organisms, complicating identification of the gene or genes that are causally associated with pigmentation variation [11, 67].

There are several limitations to our study, foremost being our modest sample size of 162 individuals, with current eQTL datasets reaching sample sizes an order of magnitude larger [49]. Many of the populations participating in this study live at considerable distances from medical or scientific facilities, and all necessary tools and supplies must be transported to field sites, greatly limiting the capacity for sample collection. Additionally, we are limited to studying blood tissues among these populations. Generation of induced pluripotent stem cells (iPSC) may allow for the study of gene regulation across developing tissues or differentiated cells within diverse populations [68, 69], but such approaches remain technically difficult. This study is also restricted to steady state gene expression, which may miss cell-type- or dynamic, environment-specific genetic effects, which cannot be captured in bulk and/or steady-state tissues [29,70,14,13,71,72]. Despite these limitations, this study makes important contributions to our understanding of gene expression variation and the molecular basis of human adaptation in sub-Saharan Africa.

## Conclusion

We have presented a comprehensive analysis of transcriptomic variation in a cohort of previously unstudied indigenous sub-Saharan Africans. We identify extensive novel regulatory variation in Africans and show that the study of African populations improves the detection of transcriptomic QTLs and fine mapping of causal variation. Studying diverse populations within Africa also allows for the detection of genes targeted by population-specific selection, including a evidence of selection on *TMEM216* expression in the Mursi and strong colocalization between *TMEM216* eQTLs and a pigmentation GWAS locus.

## Methods

### Sample Collection

Phenotypic, genealogical, and biological data were collected from individuals belonging to nine populations in Ethiopia and Tanzania. Prior to sample collection, IRB approval for this project was obtained from the University of Pennsylvania. Written informed consent was obtained from all participants and research/ethics approval and permits were obtained from the following institutions prior to sample collection: the University of Addis Ababa and the Federal Democratic Republic of Ethiopia Ministry of Science and Technology National Health Research Ethics Review Committee; COSTECH, NIMR and Muhimbili University of Health and Allied Sciences in Dar es Salaam, Tanzania. To obtain DNA and RNA data, whole blood was collected using vacutainers and RNA was stabilized in the field using LeukoLOCK Total RNA Isolation System (Ambion life Technologies). The Poly(A)Purist Kit (Ambion Life Technologies, CA) was used for mRNA selection, and Ampure XP magnetic beads (Beckman Coulter, CA) were used for size selection after amplification.

### Genotyping and imputation

A subset 162 individuals were genotyped as part of the 5M dataset using the Illumina Omni5M SNP array, which includes approximately 4.5 million SNPs. The full 5M dataset was phased using Beagle 4.0 [73] and the 1kGP reference panel [19]. These data were further imputed using minimac3 [74] and a reference panel consisting of the 1kGP and 180 WGS from the Tishkoff lab (unpublished).

### PCA and ADMIXTURE

To identify related individuals, relatedness was inferred in the imputed 5M dataset using the KING extension of plink 2.0 [75]. To place the genetic variation in this study within a global context, the 5M imputed dataset was merged with the 1KGP. Individuals from the 162 in this study with inferred relatedness more distant than third degree were then extracted from the merged dataset (145 total), along with 20 individuals each from the YRI, CEU, and CHB populations, restricting to unambiguous SNPs (i.e. excluding A/T and C/G) with MAF > 0.01 and with imputation accuracy (r^2^) greater than 0.99 reported from minimac3. SNPs were LD-pruned using plink v1.90 [76] and parameters ‘--indep-pairwise 50 10 0.1’. PCA was performed on this dataset using smartpca from EIGENSOFT v6.1.4 [77], with ‘numoutlieriter’ set to 0. ADMIXTURE [78] was run on the same dataset using parameters ‘--cv -j8 -B100 -s7’.

### mRNA sequencing and molecular trait quantification

Samples were sequenced on an Illumina HiSeq to a median depth of 56,122,076 reads (11,727,716 min., 228,660,534 max.). Prior to mapping, all reads aligned to rRNA genes with BLAST [79] were removed. Remaining reads were mapped to the hg19 genome with STAR v2.5.3a [80] and the GTEx GENCODE v19 gene annotations [81] using two-pass mapping. Expression was quantified at the gene level using featureCounts v1.5.3 [82] as fragments per gene, as well as using RSEM v1.2.31 [83] as transcripts per million (TPM). Splicing was quantified using leafcutter [84] as fraction of intron exclusion reads per cluster (JPC).

### Cell-type inference

Cell type fractions for each individual were inferred using CIBERSORT [24]. The LM22 signature gene file from Abbas *et al*. [85] was used to infer frequencies of 22 immune cell types for a mixture file of TPM values for all 171 individuals with RNA-seq data. Quantile-normalization was disabled and 1000 permutations were used.

### Quantile normalization and hidden factor inference

Prior to hidden factor inference and QTL mapping, molecular phenotype matrices were first filtered and quantile-normalized. For eQTL mapping, only lncRNA and protein-coding genes with more than 5 reads in at least 20 individuals and with mean TPM > 0.1 across all populations were considered. For sQTL mapping, introns from lncRNA and protein-coding genes with no more than 5 individuals with 0 reads were included. Furthermore, clusters were required to have at least 20 reads in at least 100 individuals and have 0 reads in fewer than 10 individuals. These filtered phenotype matrices (TPM for eQTL mapping and JPC for sQTL) were then quantile normalized using the two-stage procedure implemented by GTEx [28]. Briefly, the distribution of the phenotypes per individual were first quantile normalized to the mean of the phenotypes across individuals. Next, the distribution of each phenotype was quantile normalized to the standard normal. Hidden covariates were inferred using *PEER* [23] for these quantile-normalized phenotype matrices.

### eQTL and sQTL mapping

Expression and splicing quantitative trait loci were mapped using a linear mixed modelling approach, using the quantile-normalized gene or intron fractions as phenotypes, while correcting for sex, age, cell-type composition, delivery date, latent *PEER* factors, and genetic relatedness. Mapping was performed for SNPs with MAF > 0.05, imputation r^2^ > 0.3, and within 100kb of the target phenotype (gene TSS for eQTLs and intron for sQTLs) using *GEMMA* [26] and a genetic relatedness matrix (GRM) generated from all biallelic SNPs across the imputed, 162 individual genotype dataset. tQTL mapping was repeated across a range of *PEER* factors: 0-5, 10, 15, 20, 25, and 30 factors for eQTL mapping, and 0-10 factors for sQTL mapping, and the number of factors maximizing the number of eQTLs or sQTLs discovered were chosen for downstream analysis.

To identify significant QTLs, tested SNPs for each phenotype were first FDR corrected using Benjamini-Hochberg (BH), yielding single-corrected p-values (*P’*) for each tested SNP-phenotype pair. The minimum *P’* per phenotype were again FDR-corrected using BH, yielding double-corrected p-values (*P’’*) per phenotype, and phenotypes with *P’’* < 0.05 were considered significant. To identify significant SNPs, a threshold was set equal to the lowest *P’* for the phenotype with highest significant *P’’*, and all SNPs with *P’* lower than this threshold were deemed significant.

### Credible Sets

For each gene or intron of interest, Approximate Bayes Factors were calculated for each tested SNP using the function ‘approx.bf.estimates’ from the coloc package [86], or the function ‘approx.bf.p’ in cases where effect size or standard error information was not available. The posterior probability of each SNP *n* being causal (*PP_n_*) was then taken as:

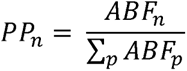

Similar to The Wellcome Trust Case Control Consortium *et al.* [87], where *ABF_n_* is the Approximate Bayes Factor of SNP *n*, and *p* indexes all tested SNPs for a given feature of interest. A 90% credible set was then defined as the minimal number of SNPs whose sum of posterior probabilities was > 0.9.

### Functional Enrichment

All SNPs in the imputed genotype dataset of 162 individuals were annotated for functional consequences using the Variant Effect Predictor (VEP) [88] with parameters ‘--per_gene --most_severe’. In addition, SNPs were overlapped with 15 state ChromHMM tracks for PBMCs (E062) from the Roadmap Epigenomics Consortium [67], transcription factor binding sites for lymphoblastoid cell lines (LCLs, GM12878) from ENCODE[34], and chromatin QTLs from Tehranchi *et al*. [33]. To test for enrichment, each FDR-significant eQTL or sQTL was matched on MAF and distance to nearest TSS or intron boundary, respectively, and the log-ratio of tQTL SNPs to matched background SNPs overlapping each functional category was taken as an enrichment score. This was repeated 10,000 times, producing an empirical distribution of enrichment scores for each functional category.

### Replication with GTEx v8

All SNPs and intron boundaries were converted to hg38 coordinates using liftOver [89]. For eQTLs, those hg19 SNPs that successfully mapped to locations in hg38 (81,928/82,144) and genes with Ensembl IDs shared between GENCODE v19 and GENCODE v26 (1,291/1,330) were considered (96,903/99,685 of possible eQTLs). Of these, 77,238 eQTLs were tested in GTEx v8 and could be compared. For sQTLs, SNPs and Ensembl IDs were required to successfully map between versions (49,706/49,794 and 772/776, respectively), and intron boundaries were required to map between GENCODE versions (738/1,118). Of these, 55,046 sQTLs were tested in GTEx. The fraction of true positives for successfully mapped tQTLs in GTEx, π_1_, was estimated using the R package *qvalue* [90].

### Conditional tQTL mapping

To identify tQTLs in the African cohort that are independent of GTEx v8 tQTLs, we performed eQTL and sQTL scans conditioning on independent GTEx eQTLs and sQTLs identified via step-wise regression [91]. In cases where there are no significant tQTLs in GTEx we instead use the top variant per feature. To account for these variants, we residualize the quantile-normalized feature matrices used in the original QTL mapping against the genotypes of independent GTEx QTLs. We then perform identical eQTL and sQTL scans, and consider genes and introns with variants that pass the original FDR threshold as independent.

### LD variation across populations

To compare LD structure between East Africans and Europeans at tQTL loci, LD was estimated (using *r^2^*) between lead SNPs for eQTLs and sQTLs and all tested SNPs in the East African and 1kGP EUR samples, restricting to those variants polymorphic in both, resulting in an *r^2^* vector per group (East Africans and EUR) per locus (eGenes and sIntrons). For each tQTL locus, we estimated the Pearson correlation *ρ* between the East African and EUR *r^2^* vectors, and the distribution of these *ρ* values was compared for tQTLs shared between East Africans and GTEx and independent tQTLs.

### eQTL mapping in 162 European-Americans from GTEx v8

eQTL mapping was performed on 162 individuals of European ancestry from GTEx v8 using FastQTL [31] with 10,000 permutations for all SNPs with MAF > 0.05 and within 100kb of the target TSS. Covariates included the top 15 *PEER* factors, top 5 genotype PCs, sex, platform, and PCR batch. Significance was evaluated using the hierarchical Benjamini-Hochberg procedure used for African samples.

### Scans of selection

To test for genetic differentiation between our African dataset and Europeans, all individuals belonging to the 9 populations in our study were extracted from the full 5M dataset (664 total) and allele frequencies were combined with frequency information for EUR populations from the 1KGP, restricting to SNPs polymorphic in both datasets. *F_ST_* was estimated using the Hudson estimator [92], and SNPs within the top 99^th^ percentile (*F_ST_* > 0.36) were considered outliers. To test for overall enrichment of *F_ST_* outliers among tQTLs, we use an approach similar to Quach *et al.* [13] The maximum *F_ST_* value of SNPs in LD with lead tQTL SNPs (r^2^ > 0.8) was found, and the fraction of outliers among these maximum *F_ST_* values was calculated. To generate a null expectation, each lead tSNP was matched with a random SNP, matching on MAF (bins of 0.05) and number of SNPs in LD (bins of [0], [1], [2], (2,5], (5,10], (10,20], (20,50], and >50). The maximum *F_ST_* of SNPs in LD with these matched SNPs was found, and the fraction of outliers among these matched maximum *F_ST_* SNPs calculated. This procedure was repeated 10,000 times, generating a null distribution of expected number of outlier SNPs.

To identify individual eGenes and sGenes with evidence of selection, weighted *F_ST_* scores were generated for each eGene and sIntron. For each feature of interest (gene or intron), the posterior probability of each tested SNP was calculated using the approach used to define credible sets, and for each feature a weighted *F_ST_* score was calculated as:

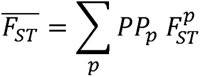

Where *PP_p_* is the posterior probability of SNP *p* being causal and 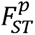 is the *F_ST_* of SNP *p*. Scores higher than the 99^th^ percentile of genome-wide *F_ST_* values were considered significant.

To detect population-specific selection, we use an adapted, polarized version of the *d* statistic for each SNP:

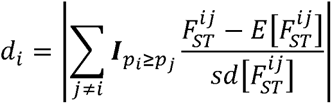

Where *p_i_* and *p_j_* are the allele frequencies in populations *i* and *j* respectively, ***I****_pi≥pj_* is an indicator function that returns 1 if *pi≥pj* and -1 otherwise, 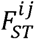 is the *F_ST_* between focal population *i* and population*j* and 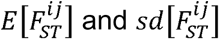 are the expected value and standard deviation of *F_ST_* between populations *i* and *j* across all SNPs. We implement this polarization procedure because SNP frequencies that are at an intermediate frequency in the focal population, but strongly differentiated in others, can show up as strong *d_i_* outliers in the focal population due to the symmetry of *F_ST_*. To identify individual eGenes and sGenes with evidence of population-specific selection, we generate weighted *d_i_* scores as described above for *F_ST_*.

Due to differential levels of admixture across populations, some *d_i_* outlier loci show genetic similarity with non-African and west-African populations, suggesting that these loci are uniquely differentiated in the focal population due to admixture. To eliminate candidates that may be driven by admixture, we also calculate the population-branch statistic (*PBS_i_*) [93] between each focal population and the CEU (a proxy for non-Africans) and the YRI (a proxy for sub-Saharan Africans):

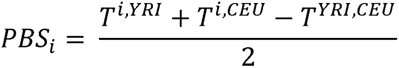

Where 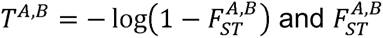 is FST calculated between populations *A* and *B* . We then go on to create a weighted *PBS_i_* statistic per gene or intron as above. Candidates of selection are then defined as those features with a *d_i_* weighted *PBS_i_* and score above the 99.5 percentile of genome-wide *d_i_* and *PBS_i_* SNP-wise statistics.

## Declarations

### Ethics approval and consent to participate

Written informed consent was obtained from all participants. IRB approval for this project was obtained from the University of Pennsylvania, and research/ethics approval and permits were obtained from the following institutions prior to sample collection: the University of Addis Ababa and the Federal Democratic Republic of Ethiopia Ministry of Science and Technology National Health Research Ethics Review Committee; COSTECH, NIMR and Muhimbili University of Health and Allied Sciences in Dar es Salaam, Tanzania.

### Authors’ contributions

SAT conceived and supervised the study. TBN, SAO, DWM, GB, WB, JBH, and AR collected and processed samples. MY and SC performed SNP genotyping. CDB, GRG, RAR, RM, and HL assisted in statistical and bioinformatic analysis. SR performed eQTL mapping of European-Americans from GTEx. DEK performed all other analyses. DEK and SAT wrote the manuscript with help from other co-authors. All authors read and approved the final manuscript.

## Acknowledgements

We would like to thank all of the study participants who make this work possible, along with our funding sources. We would also like to thank Dr. Nicholas Lahens and the ITMAT Bioinformatics Group for their assistance in data processing.

