## Supplemental Figures for "The Genetic and Evolutionary Basis of Gene Expression Variation in East Africans"

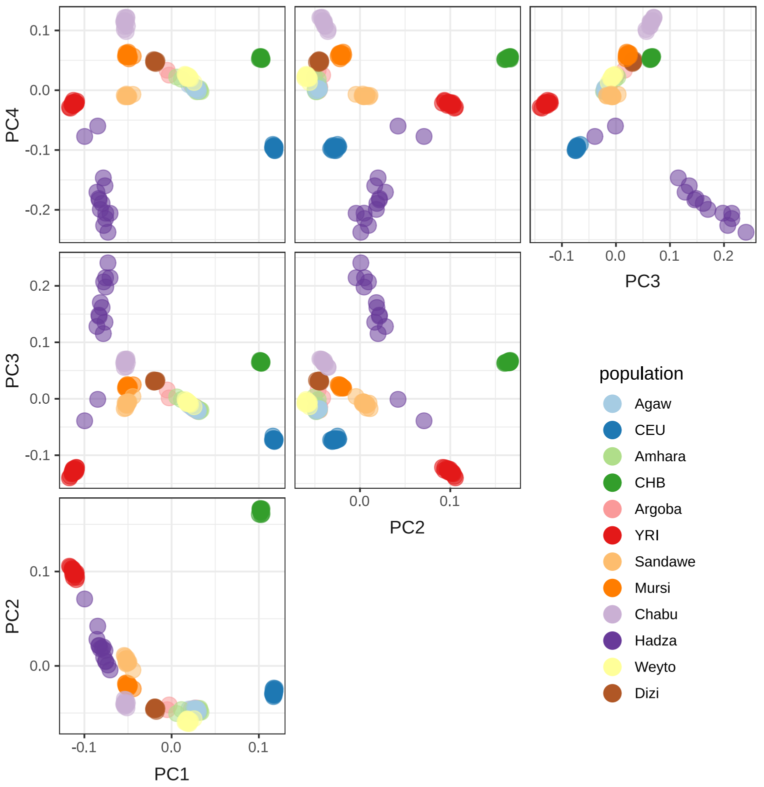


**Figure S1: Principal Component Analysis of East African and 1000 Genome Project populations**

Principal Component Analysis was performed on a merged and LD-pruned genotype dataset consisting of 145 East African individuals (filtered for relatedness) and 20 individuals each from the YRI, CEU, and CHB populations (methods). Pairwise plots of principal components (PCs) 1-4 are shown, colored by population label.

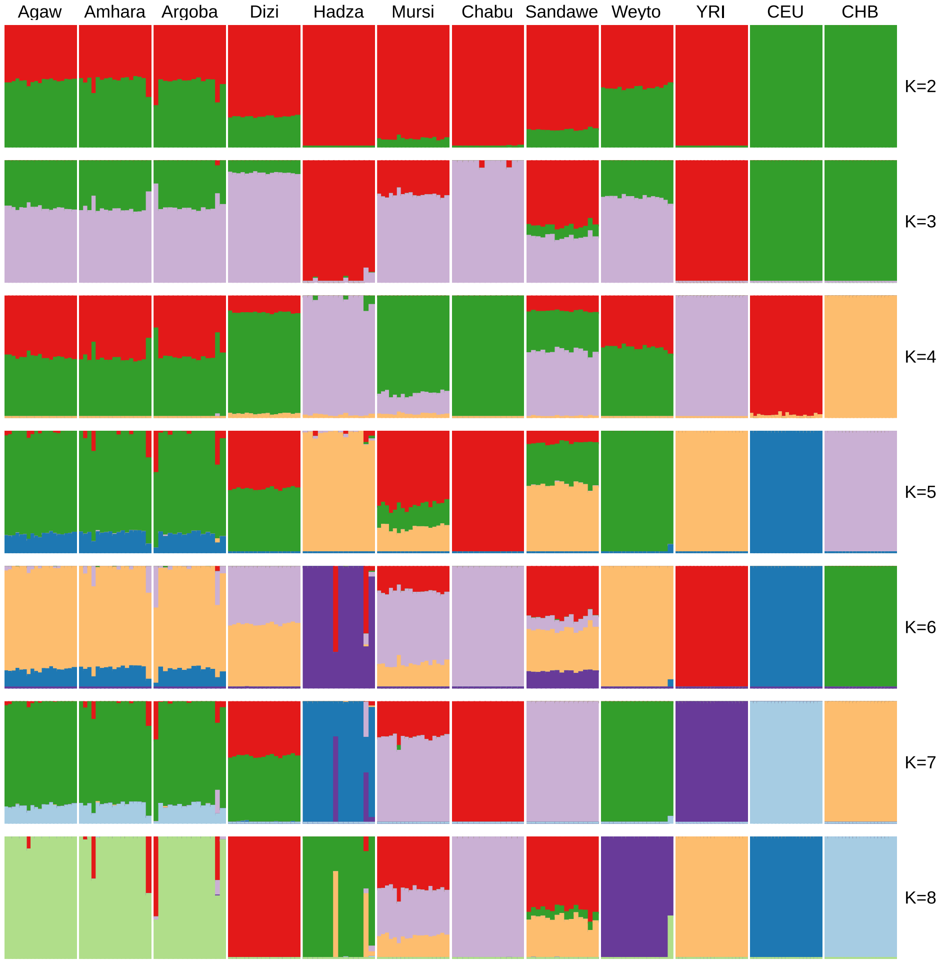


**Figure S2: ADMIXTURE analysis across K values 2-12**

ADMIXTURE analysis was performed on the merged and LD-pruned dataset used for PCA, and run for 2-8 clusters (methods). K=2 shows clear separation between Africans and non-Africans, with evidence of non-African admixture among several populations.

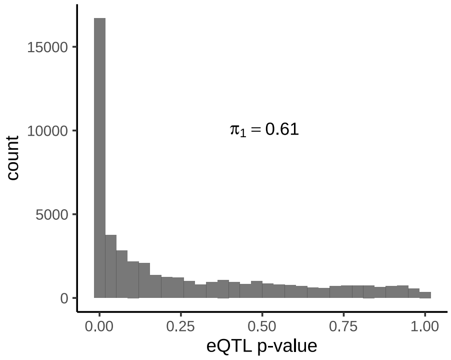


**Figure S3: $\pi_1$ of eQTL p-values of SNP-gene pairs ascertained as sQTLs**

SNP-intron pairs that meet FDR-significance are merged at the gene level to generate unique SNP-gene pairs. The $\pi_1$ is then estimated from the eQTL scan p-values of these SNP-gene pairs to approximate the number of true positives, and thus fraction of sQTLs that are also eQTLs.

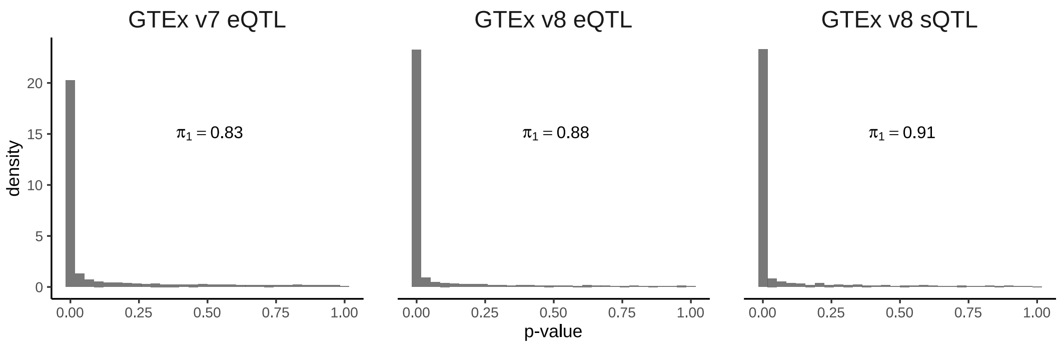


**Figure S4: π1 value of ascertained eQTLs and sQTLs in GTEx**

P-values of SNP-gene (eQTL) or SNP-intron (sQTL) pairs that meet FDR-significance in our African cohort are extracted from GTEx (v7 and v8 for eQTLs, v8 for sQTLs). $\pi_1$ is then estimated from these p-values.

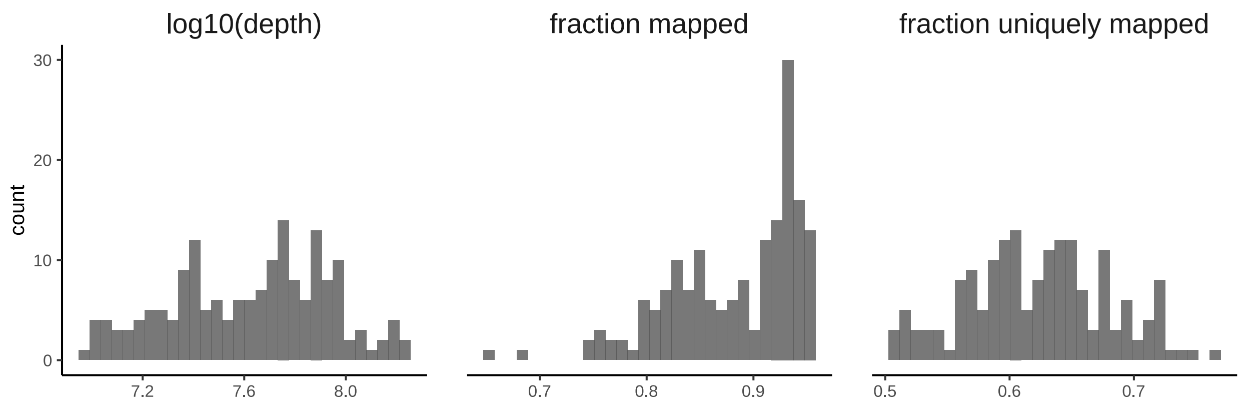


**Figure S5: Mapping statistics from STAR**

Distributions of sample mapping statistics from *STAR* (81), including *$log_{10}$* of the sample read depth (left), fraction of reads that map to the genome (center), and the fraction of uniquely mapping reads (right).

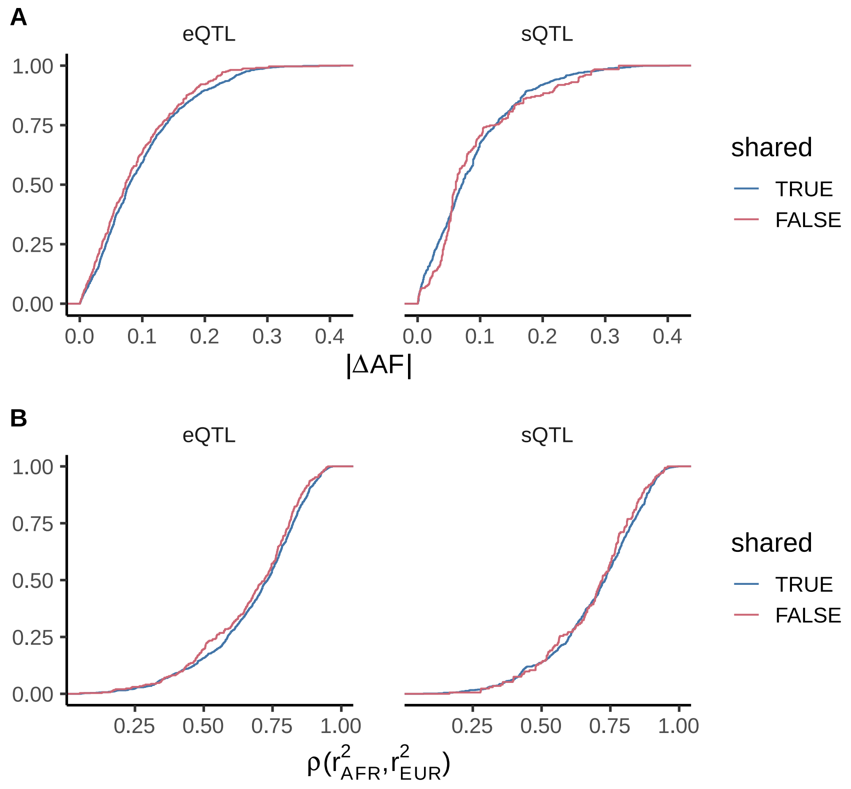


**Figure S6: Frequency and LD differences between African samples and 1000 Genomes EUR populations**

tQTL signals that do not remain FDR-significant after conditioning on independent GTEx tQTLs are coded as “shared.” **A)** The allele frequency difference of top eQTLs (left) and sQTLs (right) between our cohort and 1000 Genomes EUR populations. Independent African tQTLs are not more likely to show strong frequency differences than shared tQTLs. **B)** The correlation of African or EUR $r^2$ statistics estimated between the top eSNP or sSNP and all SNPs within 100kb. We do not find that independent African tQTLs are more likely to show weak correlations in LD.

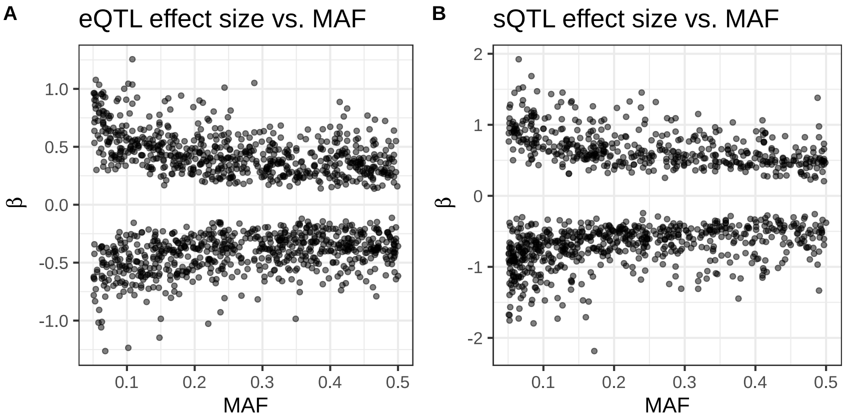


**Figure S7: tQTL effect size vs MAF**

**A)** eQTL effect sizes (β) vs minor allele frequency (MAF). **B)** sQTL β vs MAF.

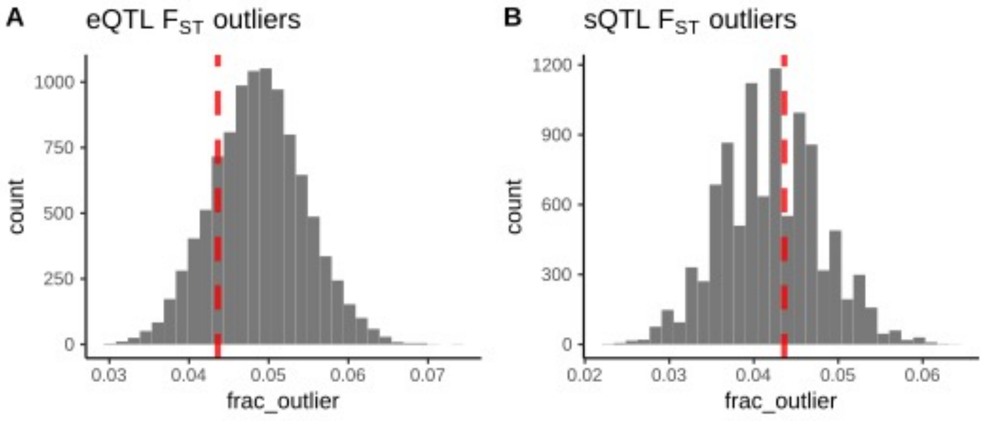


**Figure S8: Fraction of FST outliers among eQTLs and sQTLs compared with matched background**

The strongest $F_{ST}$ signal of SNPs in LD with top eQTLs (left) and sQTLs (right) is compared with SNPs matched on MAF and number of SNPs in LD. Plotted are histograms of the fraction of outlier SNP $F_{ST}$ values among 10,000 replicates (gray) and the observed fraction of outlier SNPs among the African tQTLs (red).

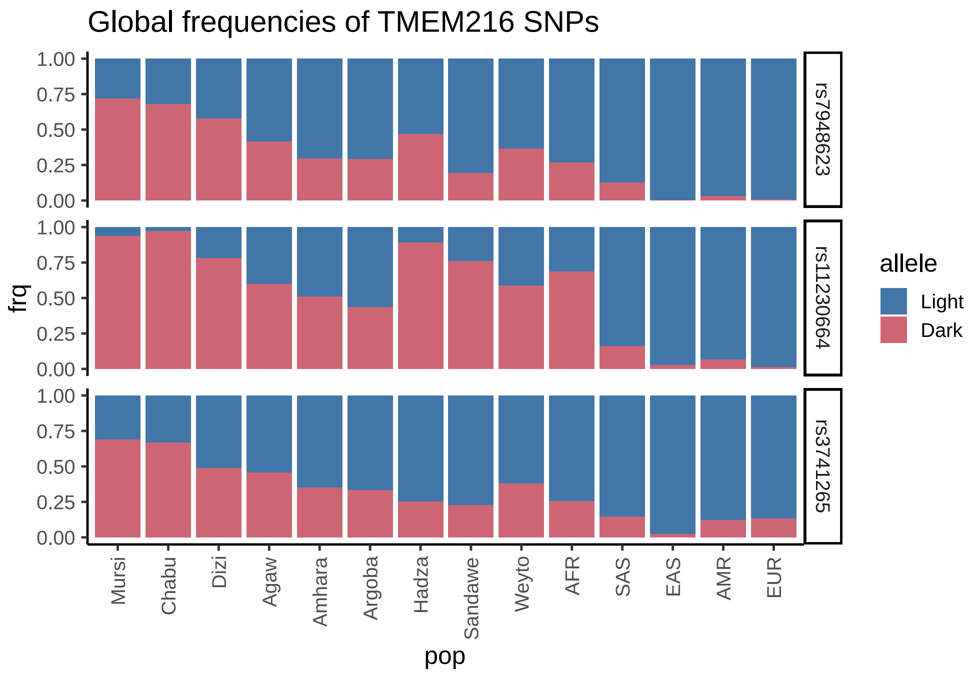


**Figure S9: Global frequencies of SNPs associated with Pigmentation variation and TMEM216 expression and splicing**

SNP frequencies for the 9 study populations and 1000 Genome Project super-populations are shown. Alleles are colored by whether they are associated with lighter (blue) or darker (red) skin pigmentation. The Mursi and Chabu show the highest dark allele frequencies at these SNPs among all global populations.

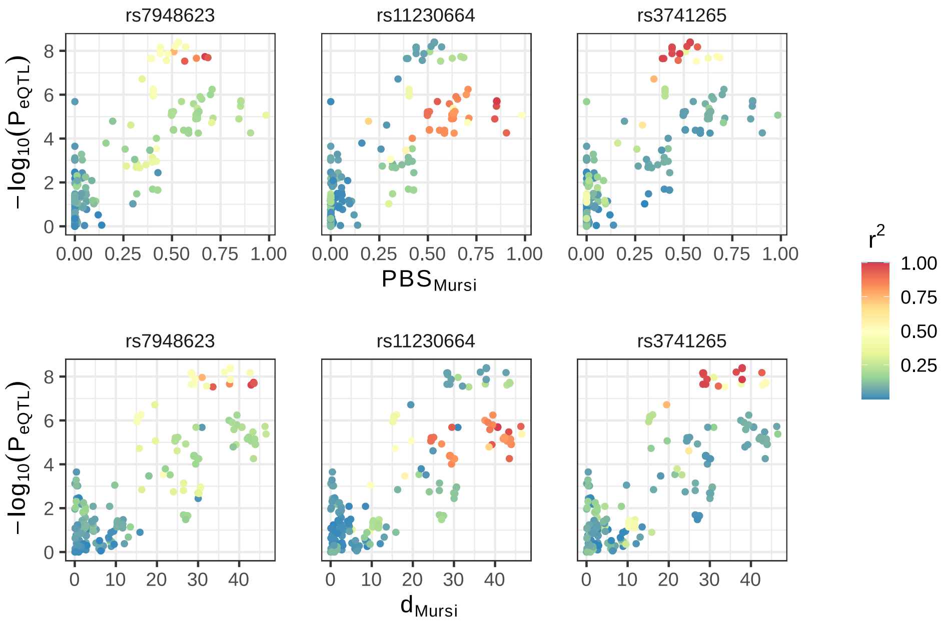


**Figure S10: Colocalization of Mursi *PBS* and *d* with *TMEM216* eQTLs**

eQTL p-values (y-axis) are plotted against Mursi *PBS* (top row) and *d* statistics (bottom row). Points are colored by their LD with the top independent pigmentation GWAS SNPs rs7948623 (left column) or rs11230664 (center column), or the sQTL rs3741265 (right column).

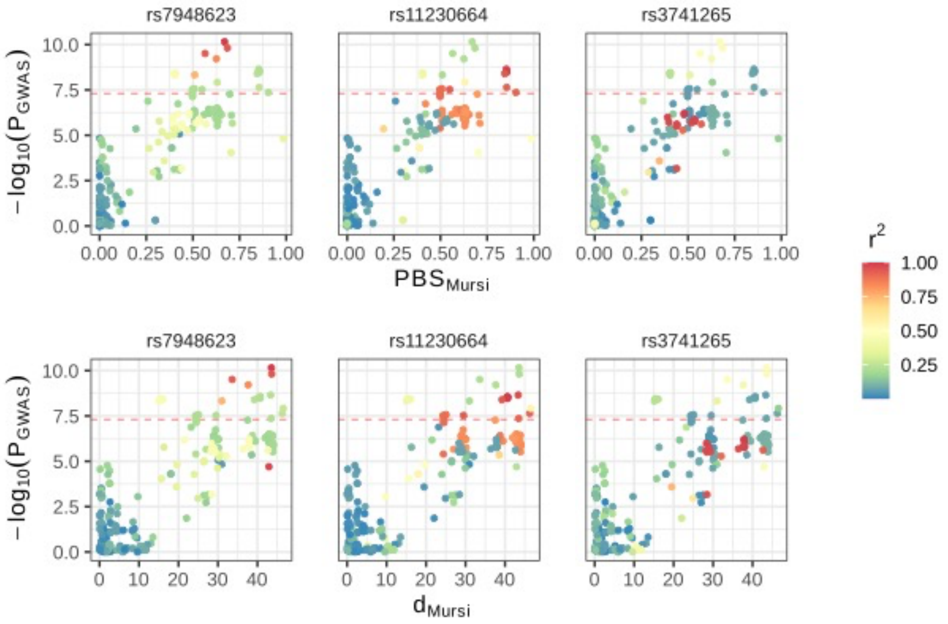


**Figure S11: Colocalization of Mursi PBS and d with pigmentation GWAS**

Pigmentation GWAS p-values (y-axis) are plotted against Mursi *PBS* (x-axis, top row) and *d* statistics (x-axis, bottom row). Points are colored by their LD with the top independent pigmentation GWAS SNPs rs7948623 (left column) or rs11230664 (center column), or the sQTL rs3741265 (right column). The red line indicates the GWAS genome-wide significance threshold of $p < 5 \times 10^{-8}$.

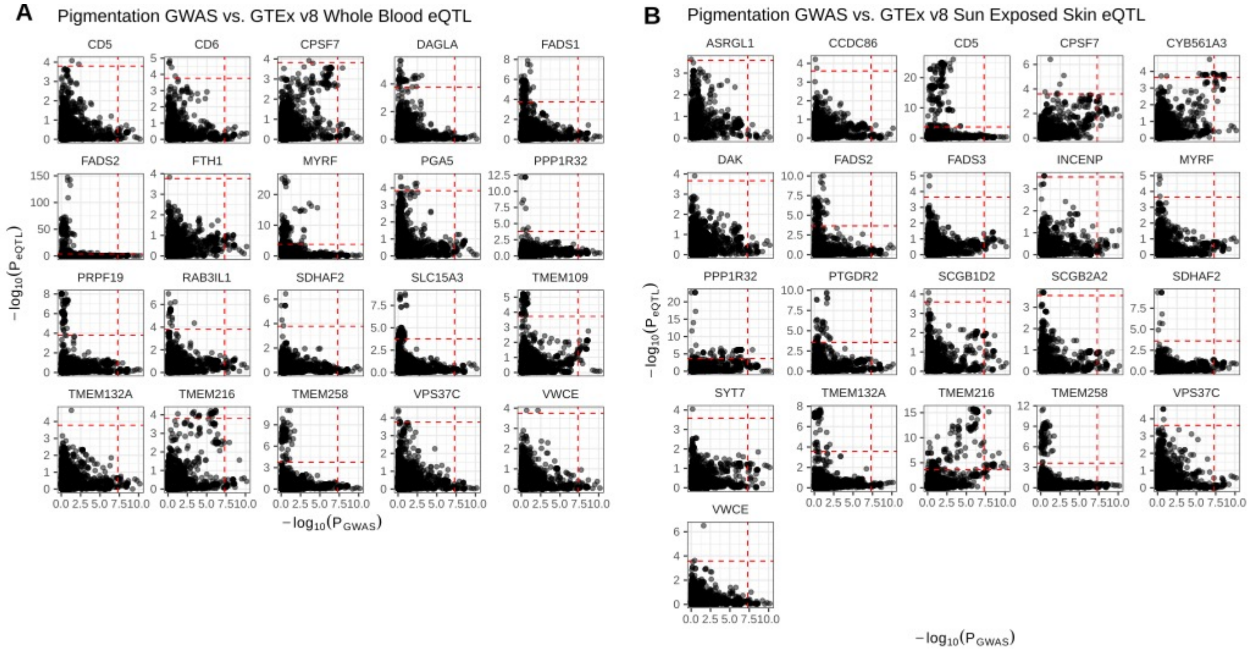


**Figure S12: 'LocusCompare' plots of African Pigmentation GWAS and GTEx v8 eQTLs**

Pigmentation GWAS p-values (x-axis) are plotted against GTEx v8 eQTL p-values (y-axis) from either (A) Whole Blood or (B) Sun Exposed Skin. Vertical dashed lines indicate the threshold for GWAS significance (5 x $10^{-8}$) and horizontal dashed lines indicate the FDR < 0.05 significance threshold for each gene.

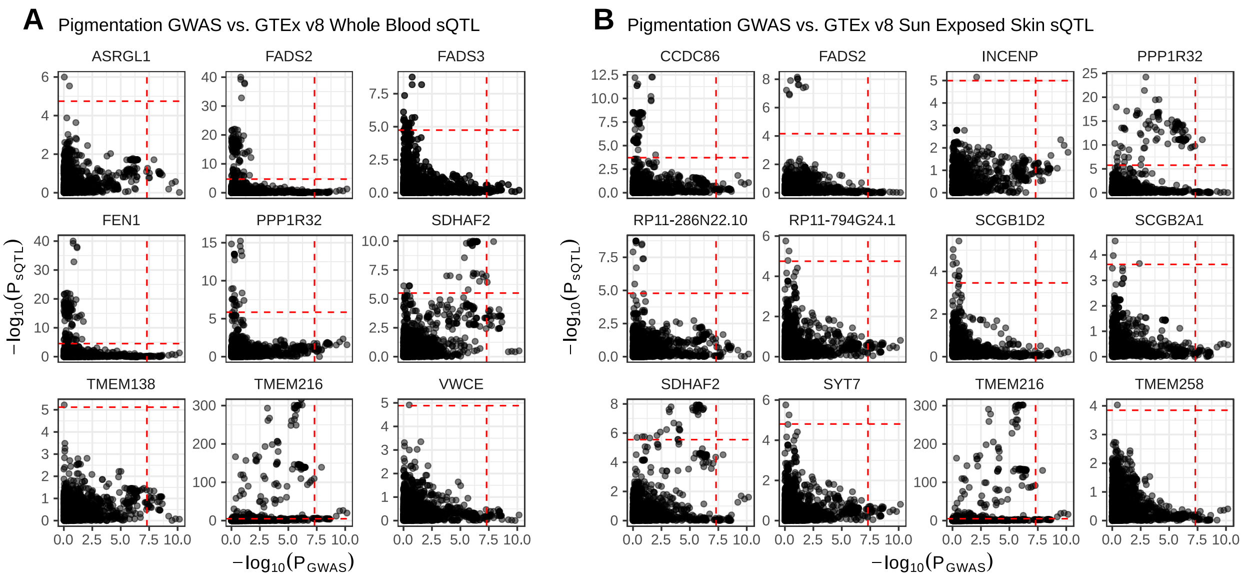


**Figure S13: 'LocusCompare' plots of African Pigmentation GWAS and GTEx v8 sQTLs**

Pigmentation GWAS p-values (x-axis) are plotted against GTEx v8 sQTL p-values (y-axis) from either (A) Whole Blood or (B) Sun Exposed Skin. Vertical dashed lines indicate the threshold for GWAS significance ($5 \times 10^{-8}$) and horizontal dashed lines indicate the FDR < 0.05 significance threshold for each gene.
